# Resolving and Quantifying Viral-Like Particles via Blind Deconvolution

**DOI:** 10.1101/2024.04.21.590467

**Authors:** Jose L Figueroa, Madeline Bellanger, Bryan Fulghum, Pieter T Visscher, Richard Allen White

## Abstract

Viruses represent the most numerous ‘biological entities’ on Earth; but the direct quantification of viruses within ecosystems reminds an ongoing challenge. The classical method of epifluorescence microscopy (EFM) reminds the gold standard measurement of viral-like particles (VLPs) within ecosystems. Quantifying VLPs in epifluorescence microscopy is burdened by ongoing challenges that include manual human counting, an absence of accurate morphological sizing, and the a range of viral sizes (20-300 nm) falling below the diffraction limit of light microscopy. Here, a proof-of-concept computer vision framework for the automated enumeration and sizing of viral-like particles is presented, known as EpiVirQuant. A novel tunable pointspread function is introduced which allows for a dynamic blind deconvolution. Final enumeration by EpiVirQuant was directly compared to manual human counting which yielded 18% more VLPs identified. EpiVirQuant quantified average VLP size of 179.5 nm, which is consistent with median size of VLPs in nature of of _160 nm. Runtime ranged from 60-80 seconds-per-image depending on parameter selection. This provides a viable proof-of-concept cost-effective solution for the enumeration and large-scale morphological analysis of VLPs.

## Introduction

Viruses represent the most abundant biological entities present, where global estimates suggest there are ∼10^31^ viral-like particles (VLPs) (1–4). Global viral abundance is larger than a mole of atoms (6.022 × 10^23^), more numerous than stars in the observable universe (∼10^21^), and greater than the number of cells in the human body (∼10^13^) (3). Quantifying VLPs within environment is critical to our understanding of the roles of viruses globally including their impact on biogeochemical cycles, lysis of biomass impacting the carbon cycle, and role in planetery health (3, 4).

Transmission electron microscopy (TEM) was originally used to determine VLPs within the environment (5, 6). TEM while still widely used to determine viral morphological characteristics failed for general use VLP enumeration. It was labor-intensive, required high cost equipment, had variability uses, and underestimated VLP abundance (7–9). Epifluorescence microscopy (EFM) provided a more costeffective, with lower cost equipment that could be transported to the field, and has been used in a wide-variety of ecosystems (e.g., aquatic, soils, and sediments) for 47 years (7–13). Throughout those 47 years of EFM many nucleic straining dyes have been used including 4’,6-diamidino-2-phenylindole (DAPI), Yo-Pro, SYBR Green I, and SYBR Gold which is utilized in this work (7–14). Epifluorescence microscopy still reminds the gold standard for viral quantification which includes high specificity, multi-color characterization, straightforward protocol, and cost-effective equipment (9).

Optical microscopy has a diffraction barrier which constrains the size of object resolution to be ∼250 nm in the lateral direction (15). Diffraction with optical defects in the lens can cause smearing, blurred spots, and can conceal resolution of fine molecular details (16). VLPs compound these issues in optical light microscopy as they are typically are sized below the diffraction limit (17). The blurring effect can be mathematically modeled as underlying process that pinpoints source to a distorted image plane when transformed this is referred to as a point-spread function (PSF) (18). While the PSF can be acquired experimental this is a challenging task in the field environment as the PSF can be sensitive to environmental conditions (19, 20). Deconvolution to a distorted image can only occur once a PSF is developed allowing for inverse transformation (21).

Here, we provide a computer vision proof-of-concept EpiVirQuant for the automated enumeration and sizing of VLPs. This proof-of-concept provides tunable point-spread function (tPSF) and a adaptive calculation for the optimal PSF. Sizing is provided by spiked-in standard fluorescent microspheres of known sizes which alleviates disparity of observed vs. true VLP size. Post blind deconvolution VLP enumeration, quantification and sizing is calculated. EpiVirQuant provides a tPSF when applications such field work or when experimental PSF is unknown (22). This is the first proof-of-concept framework that provides an opensource solution for automatic enumeration and sizing of VLPs.

## Methods

### Sample preparation

The epifluorescence microscopy were water samples prepared from Great Salt Lake (Antelope Island State Park, UT near Layton, UT 41°N 112°W) using Bellanger *et al*. method (23). Briefly samples 500 mL were filtered to 0.22 µm PVDF membrane (Millipore) and concen-trated, samples were fixed with electron-microscopy grade EM-grade glutaraldehyde until a final concentration of 2% (v/v) was attained, then flash frozen under liquid nitrogen before being stored at -80 °C until use. Samples were aliquoted into low-protein binding nuclease-free microcentrifuge tubes (ThermoFisher) with 4 µL of SYBR Gold working stock, mixed by pipette. The tubes of samples with dye were incubated on an 80 °C hot block in the dark for 15 minutes. Next, 2 µL antifade solution (10% ascorbic acid w/v in phosphatebuffered saline and filtered through a 0.22 µm PVDF membrane) was directly added to the samples, then 10 µL of samples with antifade solution were placed on slides with cover slips.

A SYBR Gold (Invitrogen S11494) working stock stain was created from concentrated dye by centrifugation at 3,500 x g for 5 minutes, then diluted at a 1:10 ratio with autoclaved and 0.22 µm molecular-grade water. The SYBR Gold working stock was then filtered using a 0.22 µm Millipore GVWP06225 PVDF (polyvinylidene fluoride) membrane then stored in darkness at -20 °C until use. Slides were prepared by cleaning with 70% ethanol then dried before soaking in 10% polylysine solution for 5 minutes. Thereafter, slides were dried in an oven at 60 °C for 1 hour. Fluorescent microspheres from the Molecular Probes’ PS-Speck Microscope Point Source Kit (P7220) with excitation and emission wavelengths of 360 nm/440 nm (blue) were added to each slide at 100x dilution (∼3.5 µL per slide). The manufacturer’s specifications list a microsphere diameter of 175 *±* 5 nm. This exceptionally small diameter makes the PS-Speck microspheres ideal as uniform, subresolution (< 200 nm), monodisperse fluorescent point sources for the calibration of instrumental optics.

The slides were imaged using an ECHO Revolve microscope with 100x oil immersion under two excitation wavelengths: *i*) FITC-channel (fluorescein isothiocyanate) green excitation light (495 nm) aimed at inducing a fluorescent response from VLPs for quantification; and *ii*) DAPI-channel blue excitation light (360 nm) which corresponds to the Molecular Probes’ PS-Speck microsphere excitation wavelength. Each EFM image was bifurcated into the corresponding DAPI (microsphere) and FITC (VLP) channels, henceforth referred to in this manuscript as the {*X*_*B*_} and {*X*_*G*_} image sets respectively. Each set of EFM images contain 20 tag image file format (tiff) 2048-by-2448 data matrices.

### Computational methods

The proposed computational methodology executes four general steps: 1) optimization box selection; 2) blind deconvolution; 3) derivation of a size-correction factor; and 4) enumeration and sizing of VLPs. A numerical value that anchors size measurements is the initial px-to-nm conversion coefficient

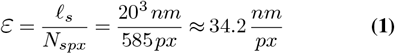

Here, *ℓ*_*s*_ is the length indicated by the scale bar and *N*_*spx*_ is the number of pixels horizontally across the scale bar which in this work were respectively 20 µm (20^3^ nm) and 585 px. In general, the *X*_*B*_ images are used to develop the tPSF and compute a size-correction coefficient supported by the known sizes of microspheres which will be expanded upon in a following section. For steps 1 and 2, an example *X*_*B*_ calibration image (GSL-01) is used for demonstration purposes, hereafter referred to as the exemplar *X*_*B*_.

### Step 1: Optimization box selection

The EpiVirQuant begins with the selection of a test pair of microspheres from the exemplar *X*_*B*_ calibration image. For this purpose, an initial coarse-object segmentation is performed using the Otsu algorithm (24). Let an image be bifurcated into two classes *{A}* and *{B}* that contain intensity values corresponding to background and foreground objects respectively. The members of *{A}* and *{B}* are separated by a latent threshold *T* which satisfies

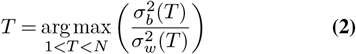

where *N* is the total number of image gray levels, with 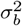 and 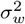 being the between-class and within-class variances respectively. After *T* is used to perform object segmentation, one morphological erosion iteration using a disk-shaped structuring element of radius 3 px was performed. This additional erosion is meant to resolve slightly doublet-shaped microsphere pairs, increasing the likelihood of a viable pair being identified. Furthermore, the optimization box is independent of size measurements, this morphological erosion is only performed for convenience within step 1. The object centroids are measured then candidate pairs within a defined proximity were retained. For this work, the proximity threshold for candidate pairs was defined as 30 px which is ≈ 1026 nm using the conversion coefficient presented in Equation 1. For the exemplar *X*_*B*_ calibration image, 4 candidate pairs were identified where the final selection was at a minimum centroid proximity of ∼449 nm. Similarly, the optimization box could be defined using a single object; however, during exploratory analyses the deconvolution effects with respect to the immediate environment were of particular interest. Furthermore, this methodology is intended for applications in substantially more crowded biological environments such as mircrobial mats in the future. Additionally, a pair of objects in the optimization box gives a distinct signature within a gray-level spatial dependence matrix as used in step 2.

The last phase of this step is to define the boundaries of the selected optimization box, *g*(*x, y*). Taking the xy limits of the microsphere pair, the bounds are extended by a padding coefficient *λ*. The applied *λ* for the exemplar case was 14 px or ~480 nm which is presented in Figure 2a. The size of the optimization box constrains the available filter sizes of the tPSF in step 2 during blind deconvolution. For example, the dimensions of the final optimization box from the examplar *X*_*B*_ were 36-by-50 px (≈ 1231-by-1710 nm). This results in a maximum tPSF filter size of 36 px, defined as *p*_*min*_. The utility of using a smaller image for deconvolution allows for the proposed PSF generator to be a rapid calculation.

### Step 2: Blind deconvolution

With the optimization box selected, the next objective is a deconvolution of the detected EFM image. This is performed absent *a priori* knowledge of the imaging equipment point-spread function (PSF) which is referred to as blind deconvolution (25, 26). The image degradation problem can be generally represented by

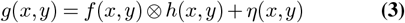

where *g*(*x, y*) is the detected noisy image, *f* (*x, y*) is the theoretical true image, *h*(*x, y*) is the point-spread function, *η*(*x, y*) represents some latent corrupting noise of the system, and ⊗ is the convolution operator. Broadly stated, the goal of deconvolution is to recover *f* (*x, y*) from *g*(*x, y*) and *h*(*x, y*).

The proposed tPSF begins with a normalized cardinal sine (sinc) function expressed using the gamma function Γ(*x*) by way of Euler’s reflection (27), shown in the one-dimensional case as

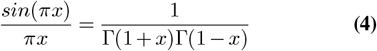

The LHS of Equation 4 is the traditional sinc function normalized by *π*, with the RHS being the gamma sinc function engrained with a *π* normalization. The gamma sinc function avoids a need for translational shifts of the tPSF arising from a point discontinuity at the origin among other convenient characteristics (28, 29). To allow flexibility during the search for an optimal solution, the gamma sinc function is parameterized by introducing two variables *τ* and *ν* that respectively act as controls for tPSF periodicity and vertical stretch. Generalizing in two dimensions, the initial tPSF is expressed as

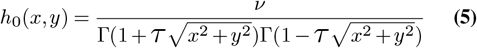

For further convenience of computation, an exponential is raised to the power of *h*_0_(*x, y*) with 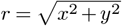

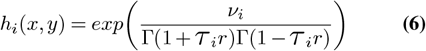

Equation 6 represents the tPSF proposed in this blind deconvolution protocol. An underlying assumption behind using a smaller optimization box for tPSF development is the deconvolution effects should scale for applications with full-size EFM images, this is highlighted in the first results section.

The next objective is to sweep over a predefined set of values for *τ* and *ν*; additionally, the filter size (*f*_*sz*_) of the tPSF is swept over. Other parameters were explored such as a leading coefficient and trailing summed term; ultimately, these had negligible effect on performance and as a result were omitted which substantially reduces the search space. The set of all optimization parameters { *f*_*sz*_, *τ, ν*} within experimentally-established ranges is defined as

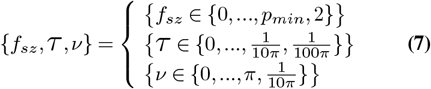

where *p*_*min*_ is the smallest odd dimension of the selected optimization box. This constraint on *p*_*min*_ is directly affected by the selected padding (*λ*) around the optimization box pair in the previous step. Since the search uses all unique com-binations of *τ* and *ν* for each *f*_*sz*_, the time complexity is *O*(*N* ^3^). This complexity can be reduced by searching the optimization landscape using more effective methods such as simulated annealing (30) or derivative-free methods (31).

When the *i*-th tPSF is constructed using the *i*-th parameter triplet, an iterative maximum likelihood estimation is performed. Here, the EFM images are assumed to originate from a Poisson process corrupted by signal-dependent noise (32). For this reason, a modified Richardson-Lucy (RL) algorithm (33, 34) is applied to the optimization box which can be succinctly shown as

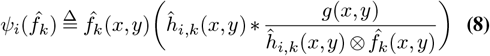

where *ψ*_*i*_(…) represents the *i*-th RL transformation of the approximated optimization box 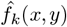 after *k* iterations. Additionally, 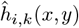 is the *k*-th tPSF estimation which was initially created using the *i*-th parameter triplet, and *g*(*x, y*) is the detected noisy optimization box as in Equation 3. The RL algorithm maintains two properties between distorted and restored images: *i*) conservation of energy; and *ii*) a positivity constraint. Maintaining the total energy is important for applications such as astronomy, electron microscopy, and EFM where the distortion does not alter the total number of photons or electrons detected (35). Furthermore, the positivity constraint is required due to the fact that photon counts at any spatial location of the detector (lens) cannot be negative.

Two metrics are introduced to quantify deconvolution performance which inform the selection of an optimal tPSF parameter triplet: information entropy and gray-level co-occurrence matrix (GLCM) energy. Given the events *S*_1_, …, *S*_*n*_ with probabilities of occurring *p*(*S*_1_), …, *p*(*S*_*n*_), the average uncertainty corresponding to each event can be discretely expressed by the Shannon entropy

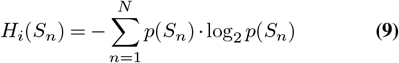

where *N* is the total number of gray levels, and *p*(*S*_*n*_) is the marginal probability distribution of the two-dimensional grayscale optimization box containing *n* events (*n* intensity values). The units for Equation 9 are bits given the base of the logarithm is 2. The minimization of Shannon entropy has been well established as a reliable metric for the quality of medical imaging and fluorescence microscopy (36).

A statistical technique for analyzing image texture which takes into consideration spatial relationships is the gray-level co-occurrence matrix (GLCM), also known as the gray-level spatial dependence matrix (37). Let (Δ*x*, Δ*y*) define a spatial offset within the image space of *g*(*x, y*) containing *q* unique intensity values. The resultant co-occurrence matrix will be of dimensions *q*-by-*q* where the (*α*-th,*β*-th) elements represent the number of times the *α*-th and *β*-th intensity values occur within the defined offset. Due to high-resolution images having a large range of pixel intensity values, a coarse-grained representation of the input image is created by grouping similar pixel intensities into a smaller number of sets. For an *n*-by-*m* image *g*(*x, y*) the *i*-th GLCM, *C*_*i*_(*α, β*), is constructed as follows

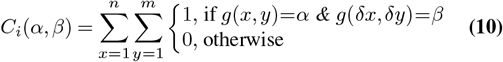

where *δx* and *δy* are respectively (*x*+Δ*x*) and (*y* +Δ*y*). The second deconvolution performance metric is GLCM energy which is a function of the *i*-th co-occurrence matrix, *E*(*C*_*i*_), defined as the square root of the angular second moment

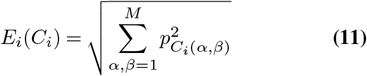

Here, *M* is the total number of gray levels and 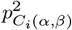 is the squared probability of the GLCM *α*-th and *β*-th intensity co-occurrence. Given the known inverse correlation of energy and entropy (38), the goal now is to maximize the quantity *E*_*i*_(*C*_*i*_) while minimizing *H*_*i*_(*S*_*n*_). Furthermore, an experimentally-balanced trade-off for *H*_*i*_(*S*_*n*_) and *E*_*i*_(*C*_*i*_) is acquired by searching for the absolute minimum difference which allows the optimization problem to be cast as

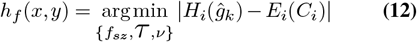

The LHS of Equation 12 is the final tPSF which is a function of the coordinates (*x, y*) provided the optimal *i*-th parameter triplet { *f*_*sz*_, *τ, ν* } . The RHS of Equation 12 represents the minimum value from the absolute difference of the optumum *i*-th entropy and GLCM energy for the estimated optimization box, *ĝ*_*k*_, after *k* iterations of the modified RL algorithm.

The first results section explores the sensitivity of solutions to Equation 12 across multiple initial conditions. Utilizing different *X*_*B*_ images for calibration generates a spectrum of image-dependent optimization landscapes that correspond to each derived tPSF. Figure 2 displays the blind deconvolution performance using the example parameter set as presented in Equation 7 which included 1887 parameter sweeps.

**Fig. 1.**
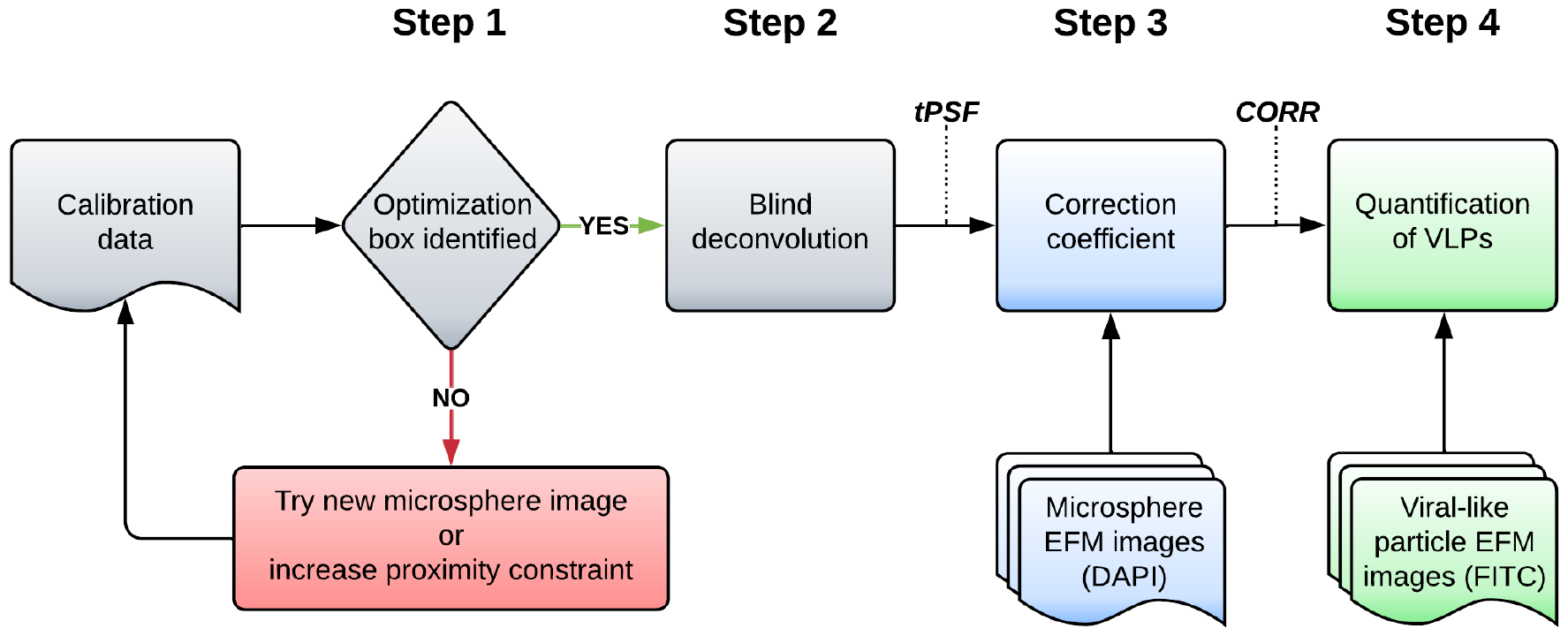
The EpiVirQuant workflow. The input to Step 1 is the selected DAPI-channel calibration image which is scanned for pairs of microspheres within a desired proximity. The microsphere pair at minimum proximity defines the optimization box; however if no pairs are identified, it is recommended to increase the proximity constraint or select a new calibration image. Step 2 performs an iterative procedure that optimizes a tunable point-spread function (tPSF) which is applied to EFM images for deconvolution. In Step 3, all available DAPI-channel EFM images are utilized to develop a size-correction factor. This scaling is then applied to the VLPs identified from each tPSF-deconvoluted FITC-channel EFM image during step 4. The measured linear eccentricities and object diameters are utilized for anomaly detection before arriving at a final VLP sizing and enumeration. Note that the selection of DAPI-channel calibration image can affect the tPSF determination, this is explored in the first results section.

**Fig. 2.**
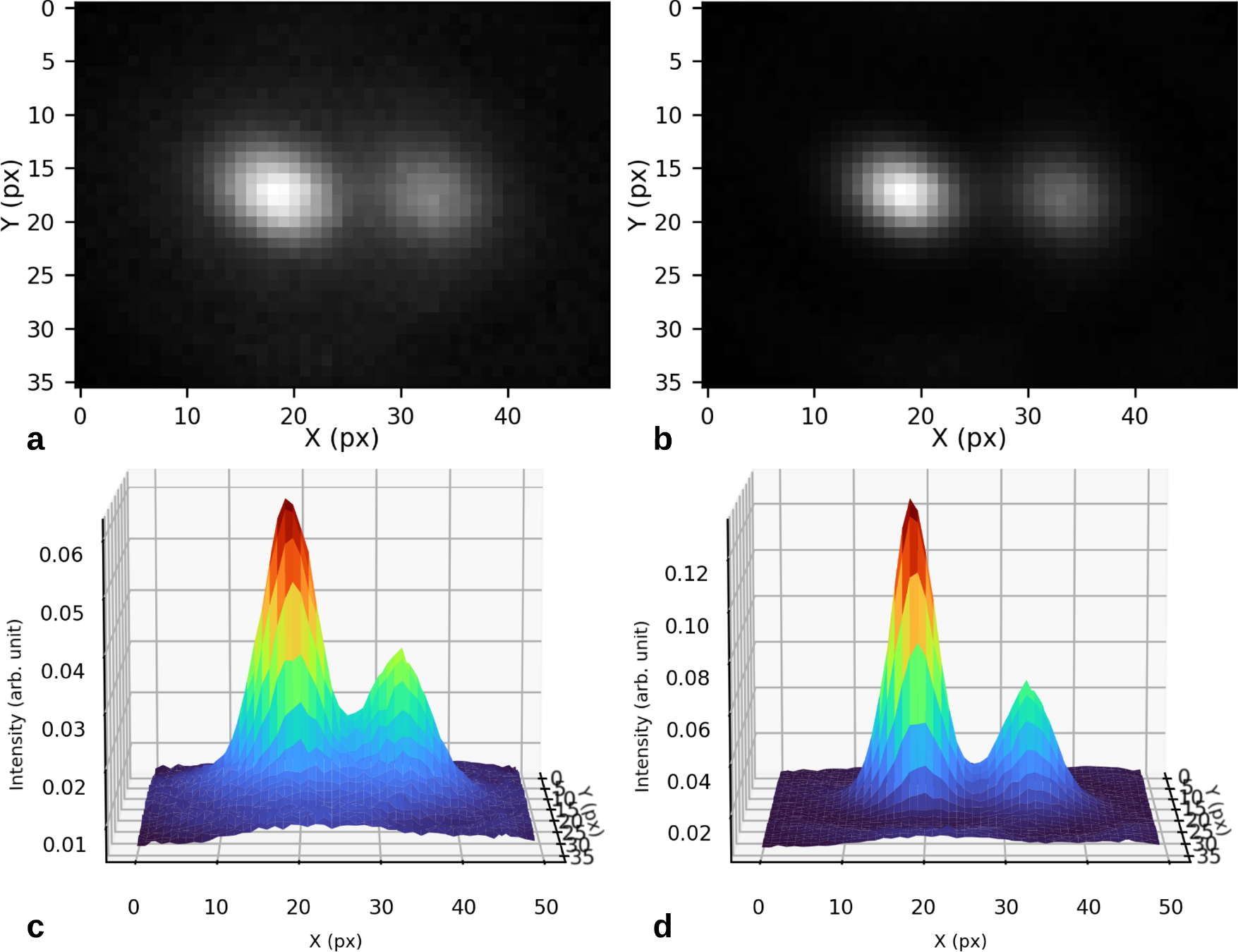
A comparison of the initial and final optimization boxes after tPSF development. (a) The initial optimization box prior to preprocessing. It is readily apparent that the fluorescence-related noise from an underlying Poisson process is present when contrasted with (b) the deconvoluted optimization box using the final tPSF. (c) The surface plot of the initial optimization box. Here, the noise-driven surface ruggedness is greater than (d), the surface of the final optimization box. Note the surrounding noise along the base is substantially decreased in (d). Additionally, the peak widths become sharper with an increased object intensity indicated by the differing surface scales.

### Step 3: Calculate size-correction factor

Equipped with the final tPSF from step 2, all available *X*_*B*_ images are utilized for the development of a size-correction coefficient, *ϱ*. This scaling coefficient is intended to correct for the discrepancy between the detected apparent object sizes and the known sizes of microspheres (*MS*_*specs*_) based on the manufacturer’s specifications, here being 175 *±* 5 nm. In this step, a statistical approximation is performed where the mean object size (*µ*_*obj*_) is calculated over all *X*_*B*_ images which allows for an informed correction scaling coefficient to be simply expressed as the ratio

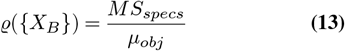

With the general objective described, step 3 begins with the application of an unmodified RL algorithm using *h*_*f*_ (*x, y*) for *N*_*RL*_ iterations ∀*X*_*B*_ in { *X*_*B*_ }. The known sizes of the microspheres allows for an optimal *N*_*RL*_ to be established. The example case from this work used the experimentally-acquired value of *N*_*RL*_ = 80. At this juncture, the additional *X*_*G*_ images can be deconvoluted using *h*_*f*_ (*x, y*) in parallel; however, to effectively measure VLPs in step 4, *ϱ* is required. As a result, only the preprocessing portion of step 4 is parallelizable within the current paradigm.

After the final deconvolution for *N*_*RL*_ iterations is complete, a per-image Otsu threshold is applied to perform object segmentation. The size measurements were taken as the mean semi-major (*a*) and semi-minor (*b*) axis lengths of the segmented objects. Therefore, for the *n*-th identified object in the resulting binary mask ∀*X*_*B*_ in { *X*_*B*_ }, the microsphere diameters, *d*(*a*_*n*_, *b*_*n*_), are measured as

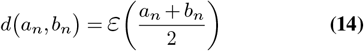

where *a*_*n*_ and *b*_*n*_ respectively correspond to the semi-major and semi-minor axis lengths of the *n*-th object, and *ε* is the px-to-nm conversion coefficient described in Equation 1. With the set of all microsphere EFM images {*X*_*B*_}, Equation 14 is used to construct the set of all segmented microsphere size measurements, given as {D}.

Next, the parametric probability density is calculated from the members of {D} shown by

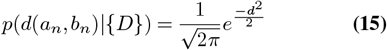

for *d*(*a*_*n*_, *b*_*n*_) ∈{*D*} which allows the *µ*_*obj*_ and standard deviation of measured objects (*σ*_*obj*_) to be calculated. Lastly, the *µ*_*obj*_ and *σ*_*obj*_ are computed ∀*X*_*B*_ in{*X*_*B*_} . This resulted in *µ*_*obj*_ = 195.04 nm and *σ*_*obj*_ = *±* 46.0 nm with the demonstration case as depicted in Figure 3. The *µ*_*obj*_ calculation was then utilized by way of Equation 13 to arrive at a final *ϱ* of 0.8973, approximately a 10.27% size reduction.

**Fig. 3.**
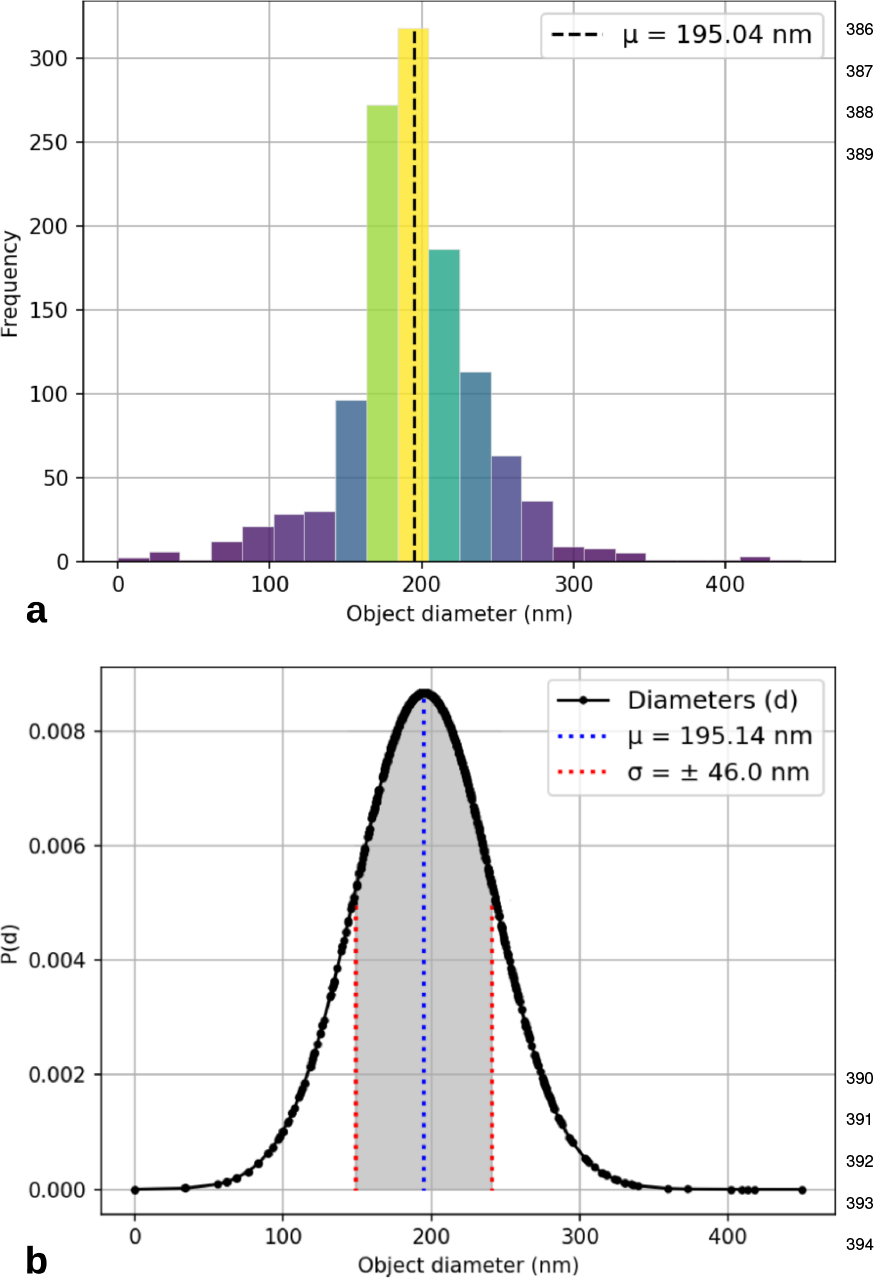
Histogram and parametric distribution of microsphere sizes. (a) The true mean microsphere size (*µ*_*obj*_) over all 20 GSL images was ∼ 195.04 nm using the exemplar GSL-01 as the calibration image. (b) The parametric density approximation allows for the measurement error to be described by the standard deviation of the distribution (*σ*_*obj*_), here being *±* 46.0 nm. There were a total of 1213 microspheres detected over 20 GSL images, resulting in ∼ 60.65 objects-per-image.

The second results section evaluates the proposed tPSF effectiveness with respect to the *µ*_*obj*_ calculation. This is exemplified by executing Steps 1-3 while varying *N*_*RL*_ from 1-300 and plotting *µ*_*obj*_ against *N*_*RL*_. It was expected that *µ*_*obj*_ should saturate around the experimentally-developed *N*_*RL*_ = 80 using the tPSF, which is further explored. The same procedure was then executed using other commonly-used PSFs to contextualize tPSF performance.

### Step 4: Enumerate and size VLPs

Similar with the previous step, Step 4 begins with the deconvolution of all *X*_*G*_ EFM images in the set {*X*_*G*_} using an unmodified RL algorithm for *N*_*RL*_ iterations with the developed tPSF from step 2. As previously mentioned, this can be performed in parallel with the deconvolution of {*X*_*B*_}. Again, a per-image Otsu threshold is derived then applied to each *X*_*G*_ ∈{*X*_*G*_} . The formerly derived *ϱ* is introduced in a modified Equation 14 to determine the final sizes of VLPs such that

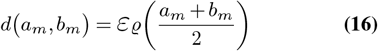

where *a*_*m*_ and *b*_*m*_ are respectively the semi-major and semiminor axis lengths of the *m*-th VLP, with *ε* and *ϱ* being the px-to-nm conversion from Equation 1 and the size-correction coefficient introduced in Equation 13 respectively.

During exploratory analysis, it was observed that further insights can be garnered from the associated object linear eccentricity (39) calculated from similar second moments as the given segmented VLPs, which is generally defined

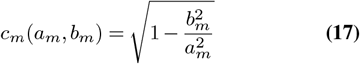

with *a*_*m*_ and *b*_*m*_ being the *m*-th semi-major and semi-minor axis lengths, assuming *a* ≥ *b*. Linear eccentricity is bound within (0,1) where 0 indicates a perfect circle. The ellipsoidal eccentricity can be used to detect false positives, or identify certain classes of microorganisms within an EFM image. For example, rod-shaped bacterium such as bacilli would have a linear eccentricity close to 1 as depicted in Figure 4.

**Fig. 4.**
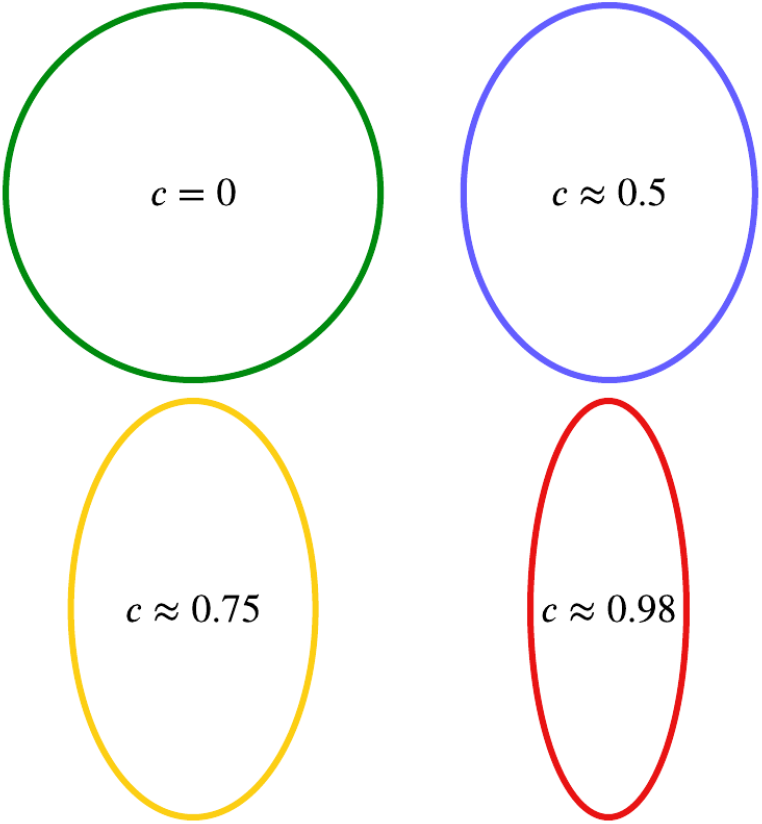
A visual for objects with varying linear eccentricities. Displayed are objects having various linear eccentricity values that may provide insights for the clustering of different biological entities. For example over many detected objects, tailed and non-tailed bacteriophages may be distinguished from one another.

A probability density is again computed by way of Equation 15 for the measured VLPs. Results from this step are presented and discussed in the last results section where EpiVirQuant enumeration is contrasted with expert human counts.

## Results

### Choice of calibration image

The sensitivity of tPSF blind deconvolution that is associated with the choice of calibration image is explored in this section. We utilized a single inital image for calibration (GSL-01.tiff) in order to establish a performance baseline due to it’s clarity and quality. Now, each microsphere EFM image is used as the calibration image (*X*_*cal*_) then the mean object size and SD are computed over the complete set {*X*_*B*_} at a fixed *N*_*RL*_ of 80 which is presented in Table 1. Note, *µ*_*obj*_ is the mean of object measurements and *σ*_*obj*_ is the standard deviation after parametric approximation which provides a quantification of measurement error; additionally, the true microsphere size was 175 *±* 5 nm.

**Table 1.** Metrics acquired utilizing different calibration images (*X*_*cal*_) represented along column 1. Columns 2-5 show the calculations over all 20 DAPI-channel EFM images for: the mean object size (*µ*_*obj*_); measurement error (*σ*_*obj*_); size-correction factor (*ϱ*); and number of total objects detected (*N*_*obj*_).

| $X_{cal}$ | $\mu_{obj}$ (nm) | $\sigma_{obj}$ (nm) | $\rho$ | $N_{obj}$ |
| --- | --- | --- | --- | --- |
| GSL-01 | 195.04 | $\pm 46.00$ | 0.8973 | 1213 |
| GSL-02 | 215.38 | $\pm 83.79$ | 0.8125 | 1333 |
| GSL-03 | 198.47 | $\pm 69.64$ | 0.8817 | 1330 |
| GSL-04 | 192.05 | $\pm 44.77$ | 0.9112 | 1223 |
| GSL-05 | 197.98 | $\pm 47.71$ | 0.8839 | 1207 |
| GSL-06 | 219.93 | $\pm 54.69$ | 0.7957 | 1216 |
| GSL-07 | 227.18 | $\pm 57.91$ | 0.7703 | 1217 |
| GSL-08 | 232.16 | $\pm 58.32$ | 0.7538 | 1220 |
| GSL-09 | 192.48 | $\pm 43.12$ | 0.9092 | 1207 |
| GSL-10 | 216.67 | $\pm 59.95$ | 0.8077 | 1236 |
| GSL-11 | 216.13 | $\pm 59.42$ | 0.8097 | 1235 |
| GSL-12 | 213.28 | $\pm 43.28$ | 0.8205 | 1204 |
| GSL-13 | 215.09 | $\pm 48.60$ | 0.8136 | 1213 |
| GSL-14 | 209.62 | $\pm 74.08$ | 0.8348 | 1304 |
| GSL-15 | 235.33 | $\pm 58.74$ | 0.7436 | 1220 |
| GSL-16 | 194.89 | $\pm 47.17$ | 0.8979 | 1215 |
| GSL-17 | 188.23 | $\pm 42.26$ | 0.9297 | 1225 |
| GSL-18 | 277.42 | $\pm 180.19$ | 0.6308 | 1767 |
| GSL-19 | 216.05 | $\pm 58.20$ | 0.8100 | 1236 |
| GSL-20 | 199.58 | $\pm 59.96$ | 0.8769 | 1265 |

The varying calculation for *µ*_*obj*_ clearly indicates that the choice of *X*_*cal*_ provides different tPSF solutions. Out of 20 total DAPI-channel microsphere images, 8 attained a *µ*_*obj*_ under 200 nm (GSL-01, GSL-03, GSL-04, GSL-05, GSL-09, GSL-16, GSL-17, and GSL-20). The *σ*_*obj*_ across all 20 images exemplifies one of the fundamental limitations of EFM, the variable excitation energy introduced to the microspheres ultimately manifests as a spectrum of observed sizes and contributes to most of the measurement error. There is an observed correlation (r = 0.79) for *µ*_*obj*_ and *σ*_*obj*_, suggesting these could be used to determine image quality when selecting an optimal calibration image. Furthermore, these results imply the choice of calibration image reduces to selecting the highest-quality image by visual inspection. There is an apparent outlier (GSL-18), having a substantially larger *µ*_*obj*_ and *σ*_*obj*_ with respect to the remaining 19 images. Although the removal of outliers is not statistically justified for the detected objects, GSL-18 can be omitted due to practical considerations. Given the measurement error is larger than the target microsphere size in this instance, the derived GSL-18 tPSF is not a viable option in practice.

The next trend involves the total number of objects detected over all DAPI-channel EFM images. This consideration was motivated by the fact that a trivial solution to the minimized *µ*_*obj*_ could involve introducing small artifacts such that the total number of objects segmented explodes, which is further displayed in the next section. Here, the total number of microspheres detected across 19/20 images (omitting GSL-18) was 1238.84 *±* 40.42 with a *µ*_*obj*_ of 209.24 *±* 14.22 nm. The top performer within {*X*_*B*_} was GSL-17 attaining a *µ*_*obj*_ of 188.23 *±* 42.26 nm resulting in a *ϱ* ≈ 0.93 (a ∼7% size reduction). It will be assumed that the tPSF derived from GSL-17 represents the PSF closest to an experimental determination. For this reason, the GSL-17 tPSF will act as the selection for applications in following sections.

### Benchmarking of tPSF against other PSFs

Here the performance of the proposed tPSF is contrasted against common PSFs used for blind deconvolution. There are 3 cases being compared: 1) tPSF; 2) Gaussian PSF; and 3) a matrix of ones normalized by the number of elements. First, a brief justification is made for the final case 2 and case 3 PSFs which will be subsequently contrasted to the tPSF from the previous section. The case 2 PSF was initially constructed at 3 filter sizes: 3-by-3, 5-by-5, and 7-by-7 each having a fixed SD of 1.0. For each filter size, the *µ*_*obj*_ was computed using the case 2 PSF ∀ *X*_*B*_ in {*X*_*B*_} with *N*_*RL*_ varying from 1-300. The top performer was selected as with the previous section which was the 5-by-5 case 2 PSF. Next, the SD was varied from 0.5-1.0 to justify the selection for the final case 2 PSF of 5-by-5 with an SD of 1.0 as presented in Figure 5a. An SD of 1.0 was selected after taking into consideration the number of objects detected. As can be seen in Figure 5a, a Gaussian 5-by-5 PSF with an SD of 0.75 appears to converge faster to the target microsphere size (∼175 nm); however, it is obvious from Figure 5b that artifacts are being introduced earlier which results in the inflation of *N*_*obj*_.

**Fig. 5.**
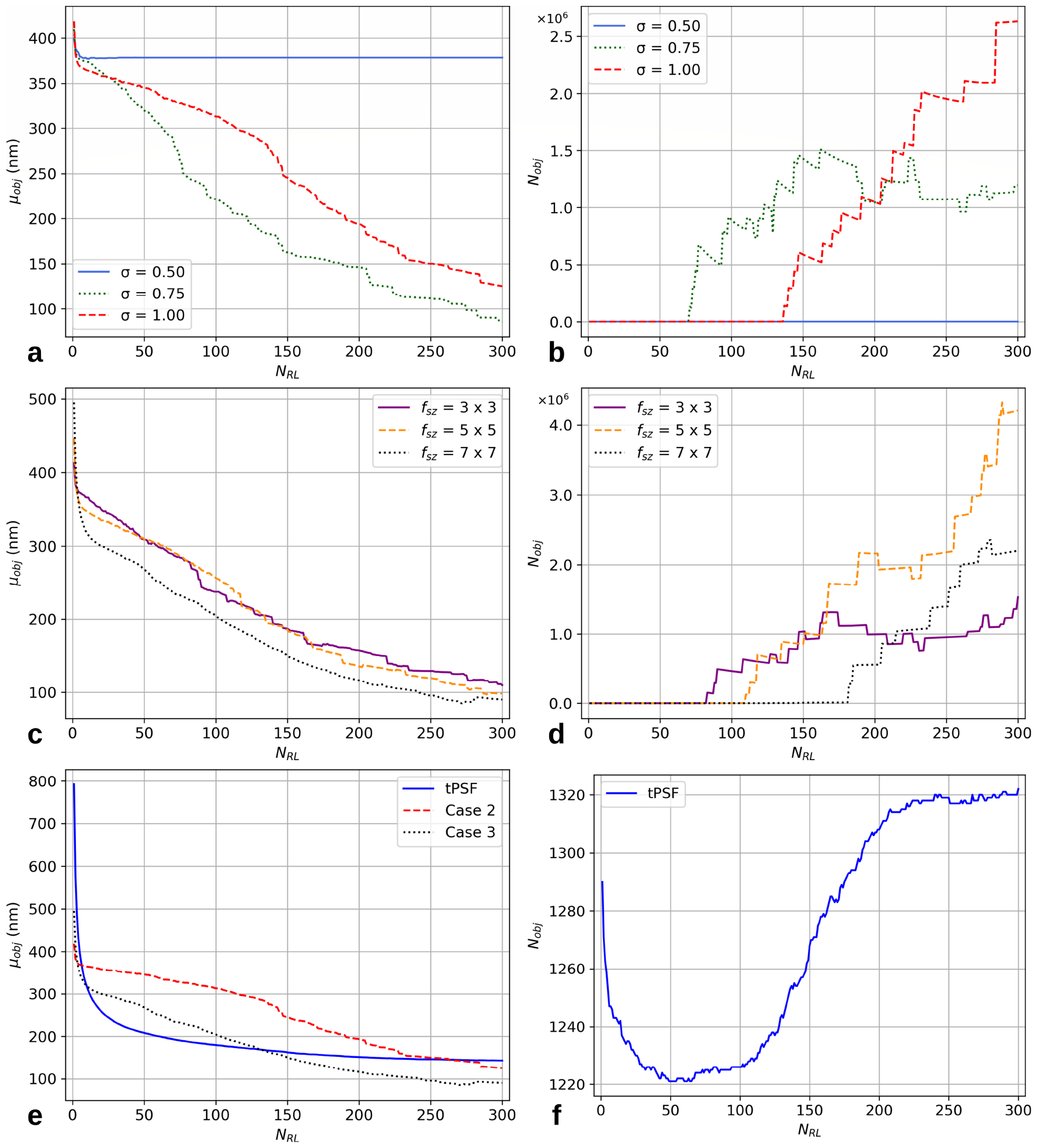
Plots of *µ*_*obj*_ vs. *N*_*RL*_ (a,c,e) and *N*_*obj*_ vs. *N*_*RL*_ (b,d,f) for case 1-3 PSFs. (a) 3 5-by-5 Gaussian PSFs with an SD of 0.5 (solid light blue), 0.75 (dotted green), and 1.0 (dashed red). The final case 2 candidate was a 5-by-5 Gaussian with an SD of 1.0 due to considerations of when each inflates to a large number of objects as shown in (b). (c) The case 3 PSF candidates at three filter sizes: 3-by-3 (solid purple), 5-by-5 (dashed orange), and 7-by-7 (dotted black). It is apparent that the case 3 PSF that converges fastest to the target microsphere size (∼ 175 nm) is the 7-by-7. This selection is further supported in (d) as the *N*_*obj*_ inflation point for this PSF occurs the latest. (e) Contrasting the case 1 tPSF to the case 2 (dashed red) and case 3 (dotted black) final candidates. It is apparent the tPSF converges to the target microsphere size in the least number of iterations. Additionally in terms of total objects detected, the tPSF is resilient to introducing artifacts as *N*_*RL*_ increases compared to the other two PSFs.

**Fig. 6.**
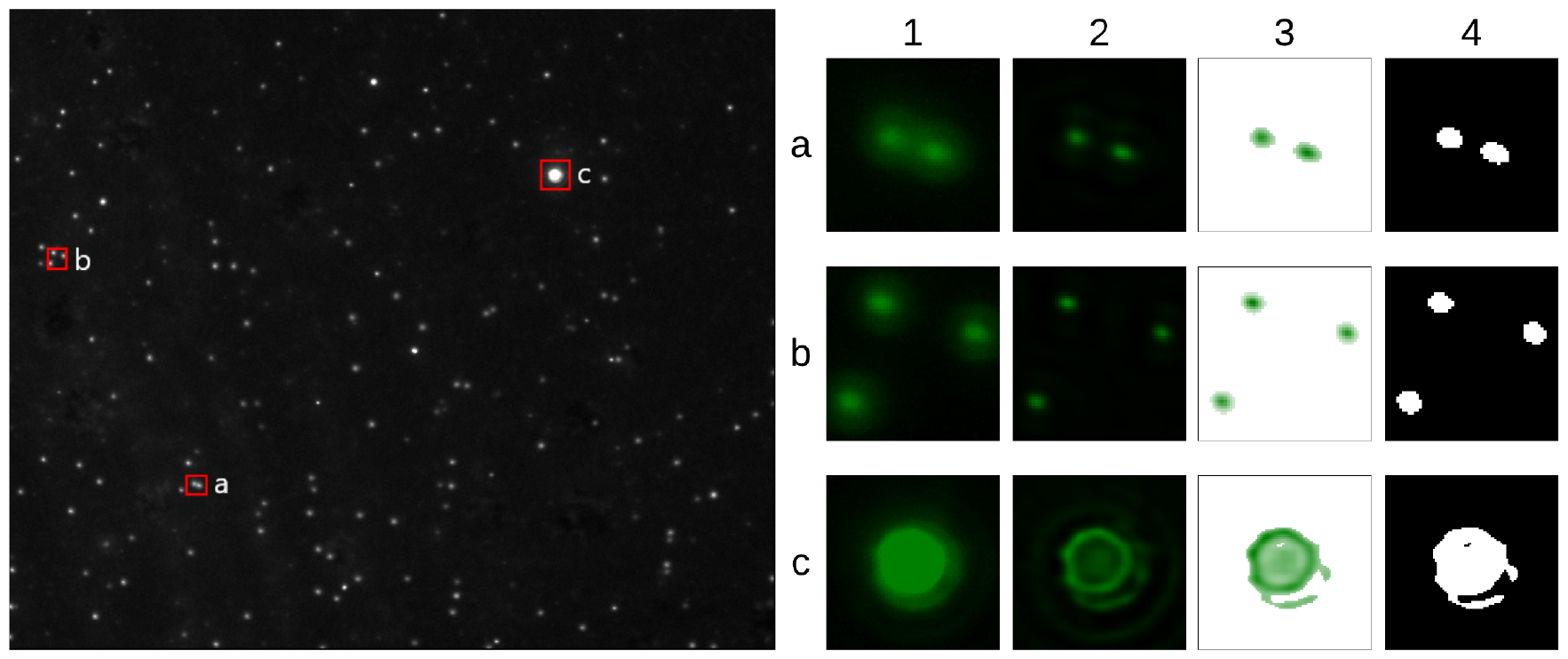
Example cases from GSL-01 (FITC) showing the (1) original objects; (2) deconvoluted objects; (3) deconvouted objects with a modified color map; and (4) binarized objects after Otsu thresholding. Three regions of interest (ROIs) are shown on the left grayscale EFM image (a-c). (a-1) The first ROI is a pair of VLPs that have overlapping Airy-disk halos which often induce false negatives when segmented. (a-2) After tPSF deconvolution, the two objects are easily resolvable. (a-3) Visualizing the deconvoluted objects using a gradient color map from white (low intensity) to green (high intensity) reveals that the underlying object is slightly larger than visible in a-2 which coincides with (a-4) the sizes of binarized objects. (b) A set of three VLPs where tPSF deconvlution reduces Airy-disk interference (b-2) providing a more accurate morphological sizing. The objects shown in (a) and (b) exemplify the large majority of objects identified by EpiVirQuant. There were 4/2012 (∼ 0.2%) objects having measured diameters greater than 500 nm such as (c). Anomalous effects can be observed after tPSF deconvolution (c-2) which should be minimized through experimental protocols.

For the final case 3 determination a similar procedure as with case 2 is performed. Again, two main considerations are taken into account: 1) which candidate case 3 PSF converges faster to the target microsphere size; and 2) which candidate has an *N*_*RL*_ that inflates latest. As shown in Figure 5c, the case 3 7-by-7 PSF converges earliest. Furthermore, the case 3 7-by-7 PSF inflates the latest as depicted in Figure 5d. The final case 2 (5-by-5; SD = 1.0) and case 3 (7-by-7) PSFs are now used to compare with the tPSF derived in the previous section. The tPSF converges to the target microsphere size earliest (*µ*_*obj*_ = 175.16 nm at *N*_*RL*_ = 114) which is exemplified in Figure 5e. At first glance, the case 3 PSF appears to compete with the tPSF in terms of quickest convergence to the target microsphere size, achieving a *µ*_*obj*_ of 175.12 nm at *N*_*RL*_ = 163; however, it reaches an *N*_*obj*_ inflation point at *N*_*RL*_ = 109 where *µ*_*obj*_ is only 246.95 nm. The observably-best performer of the three contrasted is the tPSF, both in terms of fastest convergence to the target microsphere size and with respect to the introduction of small artifacts indicated by a rapid inflation of *N*_*obj*_. Noteworthy is the stability of the tPSF to the total number of objects detected as depicted in Figure 5f. Here, *N*_*obj*_ fluctuates between 1221 (*N*_*RL*_ ≈ 50-60) and 1322 (*N*_*RL*_ = 300) microspheres whereas *N*_*obj*_ quickly inflates to a magnitude of 10^6^ for both case 2 and case 3 PSFs. It is apt to statistically infer that the likely true number of objects resides within the valley of the curve in Figure 5f around 1221-1235 microspheres.

### Enumeration and sizing of VLPs

In this Section, EpiVirQuant is utilized for the enumeration and morphological sizing of VLPs ∀ *X*_*G*_ in {*X*_*G*_} using the calibration image (GSL-17, at *N*_*RL*_ = 114) derived from the previous results sections. The first analysis involves the comparison of VLP counts made by EpiVirQuant (*C*_*c*_) and expert human counts (*C*_*h*_) as shown in Table 2. For 18/20 EFM images, EpiVirQuant identified more than human eye by an average of 19.10 *±* 15.48 VLPs. This is exemplified in column 4 of Table 2 which shows the difference between EpiVirQuant and human counts where ΔC = *C*_*c*_ - *C*_*h*_. A positive ΔC indicates that EpiVirQuant identified more than the human whereas a negative ΔC indicates EpiVirQuant underestimated the number of VLPs which occurs with GSL-12 and GSL-13. The mean size of VLPs identified was 180.90 *±* 18.32 nm which statistically falls near an established typical VLP median size of ~160 nm (17). Using the DAPI-channel GSL-17 cali-bration image does not introduce an observable bias when applied to the FITC-channel GSL-17 EFM image concerning either *µ*_*obj*_ or ΔC calculations. There were a total of 1,630 VLPs identified by human eye and 2,012 identified by EpiVirQuant. The ~19% difference was expected as computer vision techniques should identify more-nuanced low-excitation cases that are not apparent to the human eye.

**Table 2.** Enumeration of VLPs for all FITC-channel EFM images represented along column 1. Columns 2 and 3 show the VLP counts made by human eye (*C*_*h*_) and EpiVirQuant (*C*_*c*_) respectively. Column 4 is the difference between EpiVirQuant and human counts (ΔC). Column 5 shows the the mean VLP size (*µ*_*obj*_) for each image.

| $X_G$ | $C_h$ (VLPs) | $C_c$ (VLPs) | $\Delta C$ | $\mu_{obj}$ (nm) |
| --- | --- | --- | --- | --- |
| GSL-01 | 113 | 148 | 35 | 219.31 |
| GSL-02 | 117 | 133 | 16 | 170.78 |
| GSL-03 | 98 | 119 | 21 | 201.72 |
| GSL-04 | 95 | 115 | 20 | 152.30 |
| GSL-05 | 72 | 114 | 42 | 179.42 |
| GSL-06 | 102 | 123 | 21 | 156.10 |
| GSL-07 | 70 | 117 | 47 | 160.08 |
| GSL-08 | 46 | 68 | 22 | 180.47 |
| GSL-09 | 77 | 123 | 46 | 192.95 |
| GSL-10 | 104 | 120 | 16 | 165.54 |
| GSL-11 | 108 | 117 | 9 | 181.64 |
| GSL-12 | 102 | 92 | -10 | 196.00 |
| GSL-13 | 78 | 76 | -2 | 159.25 |
| GSL-14 | 75 | 85 | 10 | 197.92 |
| GSL-15 | 98 | 99 | 1 | 171.75 |
| GSL-16 | 61 | 67 | 6 | 184.91 |
| GSL-17 | 48 | 75 | 27 | 176.13 |
| GSL-18 | 66 | 74 | 8 | 168.93 |
| GSL-19 | 76 | 95 | 19 | 197.84 |
| GSL-20 | 24 | 52 | 28 | 204.91 |

An example case of GSL-01 (FITC) is presented in Figure 6 which displays two regions of interest (ROIs) representative of the majority and one large outlier. Figure 6a depicts when two VLPs are close enough that their Airy-disk halos overlap, this often induces false negatives during segmentation appearing as one large object. Figure 6a-2 shows the effectiveness of tPSF deconvolution completely resolv-ing the two VLPs, leading to a more accurate size measurement. There were a total of 4/2012 objects (∼0.2%) identified by EpiVirQuant with measured diameters > 500 nm. An instance of this is displayed in 6c with an object measured as 1290.44 nm. Given the PVDF 0.22 µm (220 nm) filter was applied during the experimental protocols, this large object may be a bacteria that made it through. Alternatively, this could be an object ~220 nm undergoing an over-excitation. In either event, it is appropriate to mitigate these cases experimentally with different filtering protocols or EFM imaging techniques.

Table 2 introduced the initial VLP measurements prior to candidate removal. Now it will be demonstrated how EpiVirQuant can be utilized for an informed VLP candidate selection protocol. All objects greater than 500 nm were omitted. Although certain viruses such as Megavirus chilensis or *Acanthamoeba polyphaga Mimivirus* can be ∼500 nm and larger (17), the clustering/classification of different virus types will be addressed in future work. Next, the linear eccentricity as defined in Equation 17 allows for a semiquantitative evaluation of the detected VLPs for the purpose of anomaly detection. Figure 7 shows a linear eccentricity vs. object diameter plot which motivates the final VLP candidate selection. There were two general cases where the linear eccentricity (c) was used for outlier removal: *i*) c > 0.975; and *ii*) c = 0. The first case was motivated by an experimentally-observed saturation point where segmented objects are indicative of rod-like structures as displayed in Figure 7b. The second case (c = 0) arises from the inherent discrete (pixelated) nature of the EFM images. A linear eccentricity of zero can be achieved in two ways, a perfect circle or a perfect square which seldomly occurs. For example, a 2-by-2 segmented object has equal semi-major and semi-minor axis lengths of ∼68.4 nm which is the resulting measurement in Figure 7d. Due to a lack of geometrical resolution for these square cases, they were omitted from the final set as a conservative measure.

**Fig. 7.**
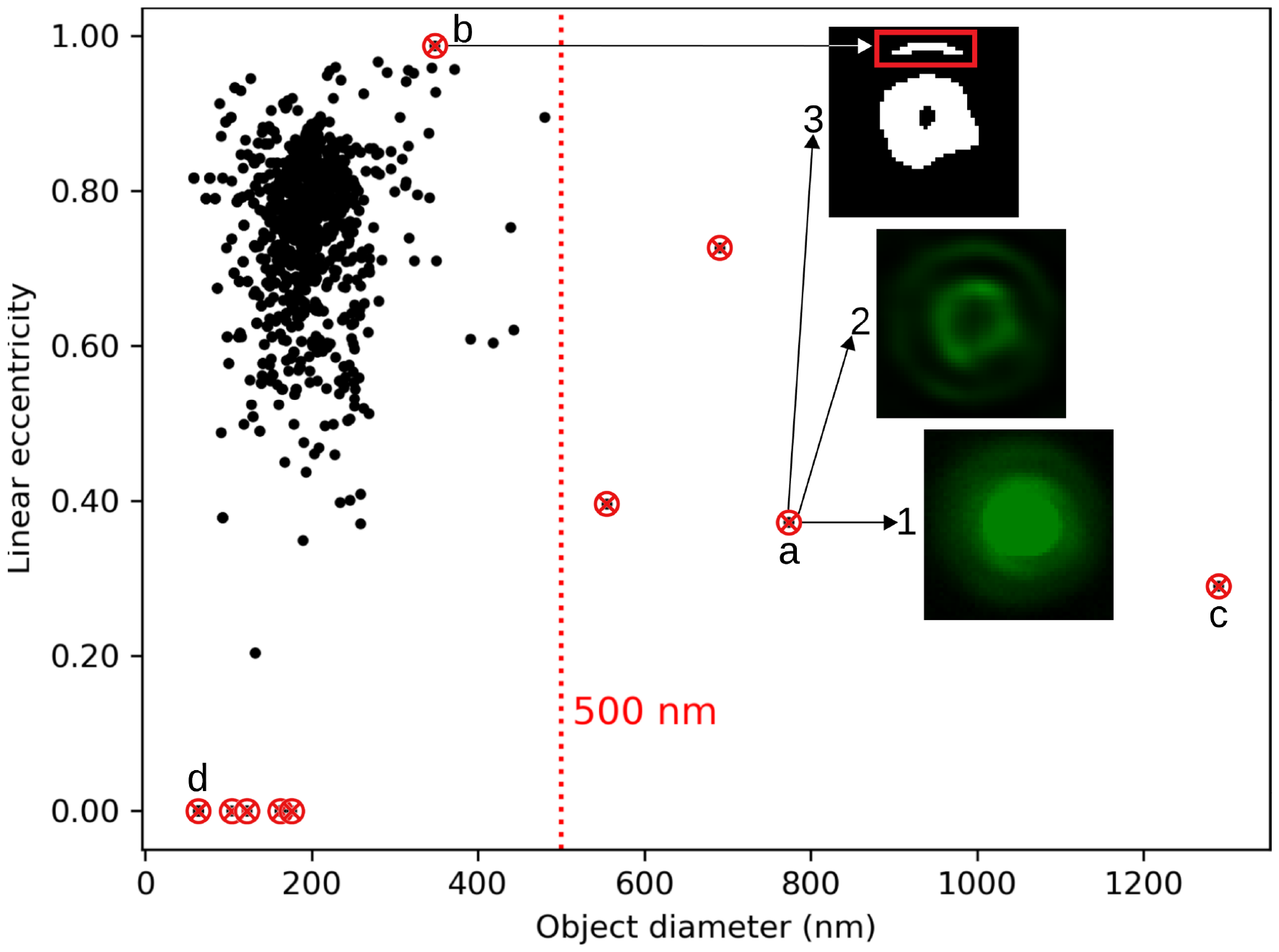
Linear eccentricity vs. object diameter plot. There were 4 objects across all 20 FITC-channel EFM images having measured diameters above 500 nm, depicted to the right of a vertical red-dashed line. A red circle with a cross indicates that the candidate object was removed from the initial set of VLPs. (a) Displayed is another example of a large bright spot with a measured diameter of 773.71 nm. This object is shown by the original (a-1), tPSF deconvoluted (a-2), and binarized (a-3) representations. (b) A long slender object was segmented in a-3 which was identified as an outlier due to a linear eccentricity of 0.9867. Although these types of cases can be mitigated experimentally, this demonstrates that EpiVirQuant has utility for the identification of artifacts through the use of linear eccentricity. (c) This object corresponds to the example introduced in Figure 6c. The final set of omitted candidates were measured to have a linear eccentricity of exactly zero. (d) Taking the object with smallest diameter (and c = 0) as an example, these instances occur when there is a symmetrical square of pixels identified as an object which results in the semi-major and semi-minor axis lengths being equal.

The final EpiVirQuant enumeration was 1,986 VLPs after candidate removal, resulting in a difference of ∼18% more objects identified compared to the human eye. There were 81/1986 (∼4.1%) objects measured less than 100 nm, 1649/1986 (∼83.0%) with sizes from 100-220 nm, and 256/1649 (∼12.9%) objects between 220 nm and 500 nm. Visualizing the VLPs using a mean intensity vs. object diameter plot as shown in Figure 8a, structure across all measurements is revealed. There exists a distinct sharp peak near the total mean object size of 179.57 nm. This experimentally can be explained by considering two factors, uniformity of staining and structural integrity of viral capsids. Essentially, the brightest VLPs (> 0.25 arbitrary units) likely represent capsids with a denser nucleic-acid packaging compared to dimmer cases. Traversing downward to the left and right of the mean, the varying sizes and intensities are explained by a combination of ruptured capsid envelopes, variable nucleic acid packaging/staining coverage, and/or cases where less excitation energy is directly incident to the capsid surface. The histogram of final VLP sizes in Figure 8b readily shows the majority of objects identified by EpiVirQuant fall within the range of ∼140 nm to ∼210 nm which corresponds to the expected sizes of VLPs previously identified in hypersaline aquatic samples.

**Fig. 8.**
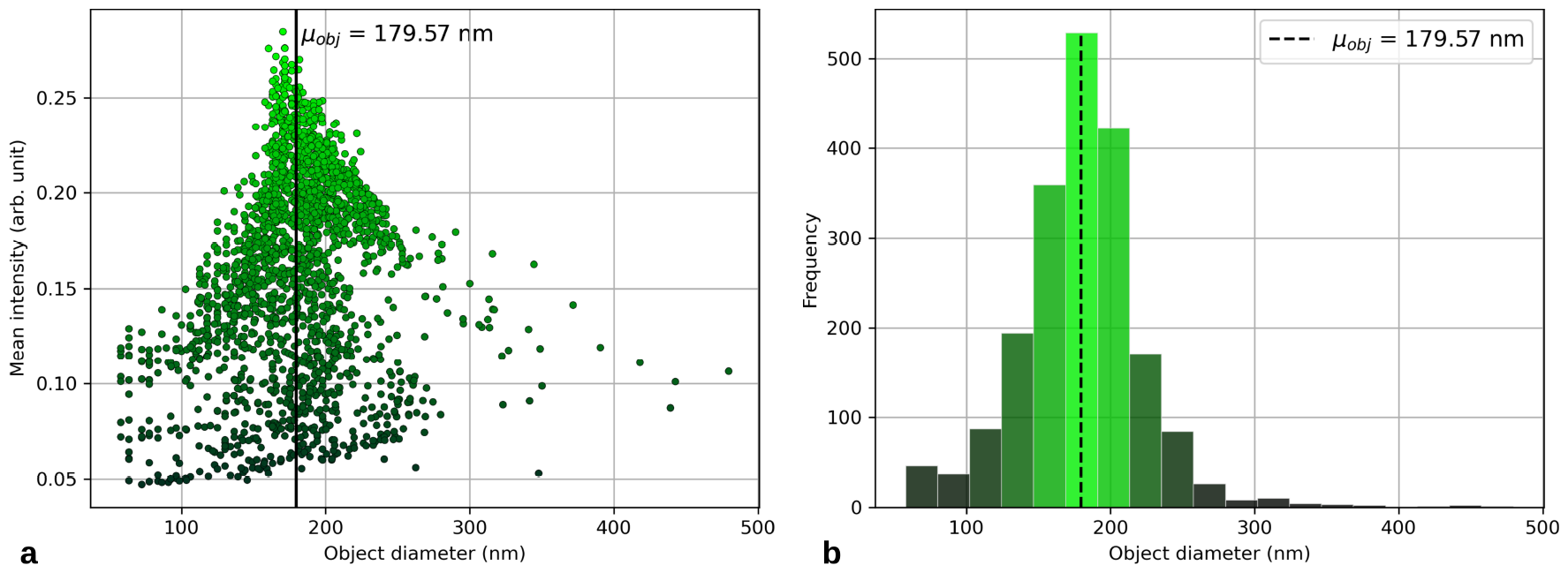
Final VLP enumeration and sizing. (a) Mean intensity vs. object diameter plot. Here, the mean intensity is calculated for all pixels that were within each segmented object. There is a clear sharp peak within intensity-diameter space which may have multiple explanations. A likely candidate explanation for the distribution in Figure 8a is an absence of perfectly uniform nucleic acid packaging and/or staining which can only be controlled to a certain degree experimentally. A solid black line indicates the mean object size over 1,986 VLPs which was 179.57 nm, this coincides with the known median size of typical viruses which is ∼160 nm. (b) Histogram of the final VLP counts. The brighter bars correspond to a larger magnitude of frequency and the dark bars indicate a smaller magnitude of object frequency within the specified range along the x-axis.

Computation times were calculated and tabulated for EpiVirQuant. At *N*_*RL*_ = 80, the complete EpiVirQuant runs in ∼60.6 seconds-per-image after an initial calibration is performed (derivation of tPSF and *ϱ*). When *N*_*RL*_ = 114, the algorithm runtime was -84.8 seconds-per-image. It can then be left to the user to decide when and how often to derive the tPSF. Computations presented in this manuscript were executed on 16 Intel Xeon Gold 6248R 3.00 GHz cores.

## Discussion

Epifluorescence microscopy has been the primary method to measure VLPs in environmental samples for longer than 40 years; yet till this day all the counting of VLPs are done by a highly trained biologist. EpiVirQuant to the best of our knowledge is the first automatic counter and sizer for VLPs currently. Here is the proof-of-concept of such a tool for VLPs within aquatic ecosystems.

We furthermore developed a novel blind deconvolution protocol which introduced a tunable point-spread function (tPSF) that harnesses both Shannon entropy and gray-level spatial dependence matrix energy for optimization which we further validated. The tPSF converges to the target microsphere size in the least number of Richardson-Lucy iterations (*N*_*RL*_ = 114) compared to commonly-used PSFs. Furthermore, the tPSF was observed resilient to the insertion of small artifacts during the deconvolution process. This was exemplified when varying *N*_*RL*_ from 1-300 where the total number of objects identified using the tPSF stabilized at *N*_*obj*_ = 1228 *±* 7 microspheres; conversely, the other PSFs quickly reach an inflation point as *N*_*obj*_ explodes to a magnitude of 10^6^. The data-driven and adaptive capabilities of the tPSF provide a new avenue for effective field applications where environmental factors are less controllable contrasted with laboratory conditions.

VLP enumeration and morphological sizing of VLPs were illuminated directly using EpiVirQuant. Using microspheres of known sizes (175 *±* 5 nm) allowed for VLP sizing which cannot be currently done with EFM VLP images this allowed for ground truth and size-correction factor (*ϱ*). The final *ϱ* developed was ∼0.93 when *X*_*cal*_ = GSL-17 (DAPI), resulting in a ∼7% size reduction for final measurements. The correlation between *µ*_*obj*_ and *σ*_*obj*_ (r = 0.79) over {*X*_*B*_} implies that the choice of calibration image from which the tPSF is derived simplifies to selecting the highest-quality image by visual inspection.

Linear eccentricity and object size were utilized for an informed candidate removal procedure. The final set of VLPs identified by the EpiVirQuant measured 1,986 VLPs compared to 1,630 identified by human eye, an increase of 18%. The cases that comprise this difference were verified as low-excitation events not visually apparent. Anomalies corresponding to the deconvolution and segmentation of larger objects (> 500 nm) were identified which should be mitigated experimentally rather than computationally. Of the 1,986 VLPs identified by EpiVirQuant, ∼4.1% had diameters less than 100 nm, ∼83.0% of diameters were measured between 100 to 220 nm, and ∼12.9% of VLPs fell within the range of 220 nm and 500 nm. The mean VLP size across all 20 FITC-channel EFM images was 179.57 nm which coincides with the typical range of ~20 nm *(Circoviruses)* to ~300 nm *(Poxviruses)*.

Many challenges still remain relating to absolute enumeration of VLPs. This includes 1) nucleic acid strand type (e.g., double vs. single stranded), 2) nucleic acid type (i.e., DNA or RNA), 3) lipid and protein content, and 4) isolation of large viruses without cellular organisms that normally are filtered out or removed (3). These problems are compounded within a mixed sample that includes mixed strand type, nucleic acid type, and the presence of cellular organisms. Advancements in super-resolution techniques may also improve VLP measurement and recovery (40).

Further modifications of the software could allow for direct enumerating of viruses within a rich-microbial environment without filtration. Future updates will include sizing for bacteria, archaea, protists, and other cellular organisms. With only minor modifications, the binary masks generated by EpiVirQuant can be directly used as training data for a deep learning framework such as convolutional neural network. This will lead to the development of robust machine predictions for quantifying VLPs after training on data for a variety of ecosystems (e.g, aquatic, terrestrial, human body, or built).

Lastly, EpiVirQuant provides a proof-of-concept sizing and enumeration for large-scale morphological analysis of VLPs, and the first step towards elucidating the global virome around us, within us, and beneath our feet in a costeffective manner.

## Data and Code availability

The epifluoresence microscopy images, scripts, beta code used in this work are publicly available at https://github.com/raw-lab/EpiVirQuant.

## Acknowledgements

We thank and acknowledge Tyler Grear for his work on this proof-of-concept. We thank Wai-Lun Lam for his work on turnable point spread function within this proof-of-concept. The authors gratefully acknowledge the University of North Carolina at Charlotte University Research Computing (URC) team that facilitated the use of high-performance computing resources during development.

## Bibliography

1. Suttle CA. Viruses in the sea. Nature, 437(7057):356–361, 2005.

2. Mushegian AR. Are there _1031 virus particles on earth, or more, or fewer? Journal of bacteriology, 202(9):e00052–20, 2020.

3. White III RA and Burns BP Visscher PT. Trends in Microbiology, 29(3):204–213, 2021. doi: 10.1016/j.tim.2020.06.004.

4. White III RA. The future of virology is synthetic. mSystems, 6(4), 2021. doi: 10.1128/msystems.00770-21.

5. Bratbak G Heldal M Bergh ØB, Yngve K. High abundance of viruses found in aquatic environments. Nature, 340(6233):467–468, 1989.

6. Heldal M Børsheim KY, Bratbak G. Enumeration and biomass estimation of planktonic bacteria and viruses by transmission electron microscopy. Applied and Environmental Microbiology, 56(2):352–356, 1990.

7. Fuhrman JA Noble RT. Use of sybr green i for rapid epifluorescence counts of marine viruses and bacteria. Aquat Microb Ecol, 14:113–118, 1998.

8. Suttle CA Weinbauer MG. Comparison of epifluorescence and transmission electron microscopy for counting viruses in natural marine waters. Aquatic Microbial Ecology, 13(3): 225–232, 1997.

9. Steele JA Schwalbach MS Hewson I Fuhrman JA Patel A, Noble RT. Virus and prokaryote enumeration from planktonic aquatic environments by epifluorescence microscopy with sybr green i. Nat Protoc, 2:269–276, 2007.

10. Fuhrman JA Proctor LM. Viral mortality of marine bacteria and cyanobacteria. Nature, 343 (6253):60–62, 1990.

11. Koike I Hara S, Terauchi K. Abundance of viruses in marine waters: assessment by epifluorescence and transmission electron microscopy. Applied and Environmental Microbiology, 57(9):2731–2734, 1991.

12. Fuhrman JA Proctor LM. Mortality of marine bacteria in response to enrichments of the virus size fraction from seawater. Marine Ecology Progress Series, pages 283–293, 1992.

13. CA Suttle. The significance of viruses to mortality in aquatic microbial communities. Microbial ecology, 28(2):237–243, 1994.

14. Fuhrman JA Suttle CA. Enumeration of virus particles in aquatic or sediment samples by epifluorescence microscopy. Manual of aquatic viral ecology, pages 145–153, 2010.

15. E Abbe. Beiträge zur theorie des mikroskops und der mikroskopischen wahrnehmung. Archiv für mikroskopische Anatomie, 9(1):413–468, 1873.

16. Corle TR Kino GS. Confocal scanning optical microscopy and related imaging systems. Academic Press, 1996.

17. Burrell CJ, Howard CR, and Murphy FA. Virion structure and composition. Fenner and White’s Medical Virology, page 27, 2017.

18. PN Dean. Confocal microscopy: principles and practices. Current protocols in cytometry, 5 (1):2–8, 1998.

19. Novak P Miks A, Novak J. Calculation of point-spread function for optical systems with finite value of numerical aperture. Optik, 118(11):537–543, 2007.

20. Jinadasa T Cole RW and Brown CM. Measuring and interpreting point spread functions to determine confocal microscope resolution and ensure quality control. Nature protocols, 6 (12):1929–1941, 2011.

21. Ramaswamy NK Satish P, Srikantaswamy M. A comprehensive review of blind deconvolution techniques for image deblurring. Traitement du Signal, 37(3), 2020.

22. Murtagh F Pantin E, Starck JL. Deconvolution and blind deconvolution in astronomy. Blind Image Deconvolution, pages 301–340, 2017.

23. Bellanger M, Visscher PT, and White III RA. Viral enumeration using cost-effective wet-mount epifluorescence microscopy for aquatic ecosystems and modern microbialites. Applied and Environmental Microbiology, 89(12):e01744–23, 2023. doi: 10.1128/aem.01744–23.

24. N Otsu. A threshold selection method from gray-level histograms. IEEE transactions on systems, man, and cybernetics, 9(1):62–66, 1979.

25. Kundur D and Hatzinakos D. Blind image deconvolution. IEEE signal processing magazine, 13(3):43–64, 1996.

26. Rameshan R Chaudhuri S Velmurugan R Chaudhuri S, Velmurugan R and Rameshan R. Blind deconvolution methods: A review. Blind Image Deconvolution: Methods and Convergence, pages 37–60, 2014.

27. Euler L. De summis serierum reciprocarum. Commentarii academiae scientiarum Petropolitanae, pages 123–134, 1740.

28. Gauss CF. Disquisitiones generales circa seriem infinitam 1+ α β/γx+⃛. Gesammelte Werke, 3:1866–1929, 1812.

29. Edwards HM. Riemann’s zeta function dover publications. New York, 2001.

30. Vecchi MP Kirkpatrick S, Gelatt Jr CD. Optimization by simulated annealing. science, 220 (4598):671–680, 1983.

31. Sahinidis NV Rios LM. Derivative-free optimization: a review of algorithms and comparison of software implementations. Journal of Global Optimization, 56:1247–1293, 2013.

32. Kingman JFC. Poisson processes, volume 3. Clarendon Press, 1992.

33. Richardson WH. Bayesian-based iterative method of image restoration. JoSA, 62(1):55–59, 1972.

34. Lucy LB. An iterative technique for the rectification of observed distributions. The astronomical journal, 79:745, 1974.

35. Conchello JA Lichtman JW. Fluorescence microscopy. Nature methods, 2(12):910–919, 2005.

36. Lee Y Tsai DY and Matsuyama E. Information entropy measure for evaluation of image quality. Journal of digital imaging, 21:338–347, 2008.

37. Shanmugam K Haralick RM and Dinstein IH. Textural features for image classification. IEEE Transactions on systems, man, and cybernetics, (6):610–621, 1973.

38. Bruillard PJ Marsh JN Wickline SA Hughes MS, McCarthy JE. Entropy vs. energy waveform processing: A comparison based on the heat equation. Entropy, 17(6):3518–3551, 2015.

39. Thomas GB. Calculus and analytic geometry. Technical report, 1968.

40. Heintzmann R Schermelleh L and Leonhardt H. A guide to super-resolution fluorescence microscopy. Journal of Cell Biology, 190(2):165–175, 2010.

